# Pharmacological CDK4/6 inhibition promotes vulnerability to lysosomotropic agents in breast cancer

**DOI:** 10.1101/2024.08.22.609150

**Authors:** Jamil Nehme, Sjors Maassen, Sara Bravaccini, Michele Zanoni, Caterina Gianni, Ugo De Giorgi, Abel Soto-Gamez, Abdullah Altulea, Teodora Gheorghe, Boshi Wang, Marco Demaria

**Affiliations:** European Research Institute for the Biology of Ageing (ERIBA), University Medical Center Groningen (UMCG), University of Groningen (RUG), Groningen, The Netherlands; IRCCS Istituto Romagnolo per lo Studio dei Tumori (IRST) “Dino Amadori”, Meldola, Italy

## Abstract

Breast cancer is a leading cause of mortality worldwide. Pharmacological inhibitors of Cyclin- Dependent Kinases (CDK) 4 and 6 (CDK4/6i) inhibit breast cancer growth by inducing a senescent-like state. However, the long-term treatment efficacy remains hindered by the development of drug resistance. Clearance of senescent-like cancer cells may extend the durability of treatment. In this study, we showed that CDK4/6i-treated breast cancer cells exhibit various senescence-associated phenotypes, but remain insensitive to common senolytic compounds. By searching for novel vulnerabilities, we identified a significantly increased lysosomal mass and altered lysosomal structure across various breast cancer cell types upon exposure to CDK4/6i in preclinical systems and clinical specimens. We demonstrated that these lysosomal alterations render breast cancer cells sensitive to lysosomotropic agents, such as L- leucyl-L-leucine methyl ester (LLOMe) and salinomycin. Importantly, sequential treatment with CDK4/6i/lysosomotropic agents effectively reduced the growth of both Hormone Receptor- positive (HR^+^) and triple-negative breast cancer (TNBC) cells in vivo. This sequential therapeutic strategy offers a promising approach to eliminate CDK4/6i-induced senescent(-like) cells, potentially reducing tumor recurrence and enhancing the overall efficacy of breast cancer therapy.

## Main text

Despite considerable improvements in early detection and treatment, cancer remains one of the leading causes of death worldwide. Breast cancer is the most prevalent cancer type among women (Wilkinson & Gathani, 2022). One of the most common anticancer strategies relies on targeting mechanisms that lead to unrestrained proliferation. This can be accomplished by inflicting high amounts of DNA damage using genotoxic stressors such as chemotherapeutic agents or ionizing radiation. However, many chemotherapeutics used to treat cancer lack specificity and their systemic administration is associated with multiple short- and long-term adverse reactions (Demaria et al, 2017; Yao et al, 2020). More targeted antiproliferative approaches, such as inhibiting Cyclin-dependent Kinases (CDK) 4 and 6 to block the cell cycle transition from G1 to S phase, have been shown to be better tolerated and have significantly fewer side effects (Wang et al, 2022). The CDK4/6 inhibitors (CDK4/6i) palbociclib, ribociclib, and abemaciclib have been approved by the FDA for the treatment of metastatic hormone receptor-positive (HR^+^) and human epidermal growth factor receptor 2-negative (HER2^-^) breast cancer (Eggersmann et al, 2019). These agents have demonstrated to provide a notable improvement in progression-free survival (PFS) compared with endocrine therapy alone (Maltoni et al, 2024; O’Leary et al, 2016; Rocca et al, 2014; Rocca et al, 2017). However, CDK4/6i are rarely cytotoxic but are more often cytostatic. Our laboratory has previously shown that treatment with CDK4/6i causes senescence-like phenotypes associated with transient or permanent growth arrest, depending on the genetic background and treatment duration (Wang & Demaria, 2021; Wang et al, 2022). Cell cycle arrest is one of the hallmarks of cellular senescence. Therefore, promotion of cancer cell senescence is a potent barrier to tumorigenesis and the desired outcome of cancer treatment (Ewald et al, 2010; Lee & Schmitt, 2019). While cancer senescence can be induced at a lower drug dose than that required for cell death, thus reducing potential toxicity, there is evidence that subpopulation(s) of senescent cancer cells might be capable of re-entering the cell cycle, acquiring stem cell-like phenotypes, and promoting disease recurrence (Evangelou et al, 2023; Schmitt et al, 2022). Therefore, a second therapy that can remove senescent-like cancer cells by targeting the vulnerability induced by the first drug, a strategy known as one-two punch, is a promising approach for maximizing therapeutic efficacy (Sieben et al, 2018; Wang & Bernards, 2018).

Lysosomes are crucial for supporting the enhanced proliferative capacity of cancer cells because of their roles in cellular metabolism, nutrient recycling, and the regulation of cellular processes vital for survival and growth (Kallunki et al, 2013). Cancer cells often exhibit alterations in lysosomal content, size, and structure (Kallunki et al, 2013). These changes render cancer cells vulnerable to lysosomal membrane permeabilization (LMP). Agents that induce LMP, also known as lysosomotropic agents, have emerged as novel therapeutic strategies for the treatment of different types of cancers (Hu & Carraway, 2020). Changes in lysosomal content and structure are also a major senescence-associated phenotype, often used for the identification of senescent cells (Hernandez-Segura et al, 2018).

In this study, we observed lysosomal changes as a prominent phenotype induced by CDK4/6i in different breast cancer cell types and evaluated the potential therapeutic effect of exposing CDK4/6i-treated breast cancer cells to lysosomotropic agents.

### Abemaciclib induces a senescent phenotype unresponsive to typical senolytics in breast cancer cells

To assess whether exposure to CDK4/6i confers senescence-like features to HR^+^ breast cancer cells, we treated MCF-7 cells with abemaciclib for 8 days. Treatment led to enlarged and flattened morphology, increased senescence-associated β-galactosidase (SA-β-Gal) activity, and diminished clonogenic capacity and EdU incorporation, indicating a senescent state (Fig 1A-C). Similar to what we have previously reported in the context of normal cell (Wang et al, 2022), treatment of MCF-7 cells with abemaciclib was accompanied by upregulation of p53, but not NF-κB, target genes (Fig EV1A). However, even after prolonged exposure to abemaciclib (30 days), signs of senescence escape were observed upon drug withdrawal, as evidenced by rescued proliferative activity (Fig 1D). This observation was in line with previous cell culture and *in vivo* reports that senescent cancer cells might escape a stable cell cycle arrest (Evangelou et al, 2023), and supports the idea that a second sequential treatment is necessary to increase therapeutic efficacy. Considering the appearance of senescence-like features, we examined the sensitivity of the abemaciclib-treated cells to various common senolytic compounds. However, abemaciclib- treated cells were insensitive to all the tested senolytics (Fig 1E). Notably, most, if not all, senolytic compounds induce cell death via the apoptotic pathway. However, cancer cells often develop mutations that render them resistant to apoptosis induction (Kulbay et al, 2022). Thus, the identification of drugs capable of triggering cell death through mechanisms independent of the apoptotic pathway could represent a more effective strategy.

**Figure 1.**
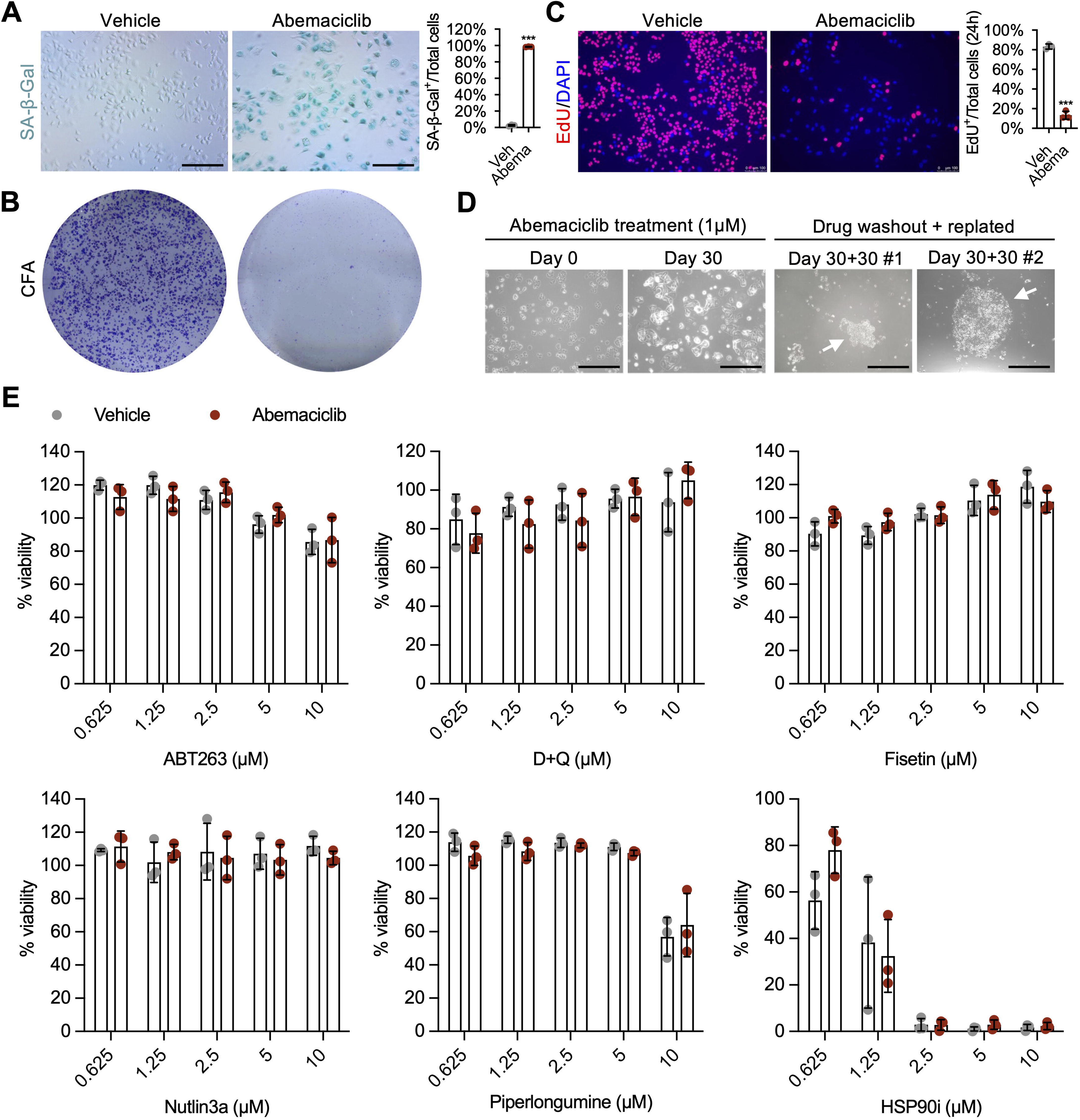
Abemaciclib-induced senescent MCF-7 cells are not sensitive to common senolytic compounds. **A,B** MCF-7 cells were treated with either vehicle (water) or abemaciclib (1μM for 8 days), then cells were replated for SA-β-Gal staining (scale bar 1mm) and quantified (**A**) or colony formation assay (for 8 days culture) (**B**), n=3 independent experiments. **C** MCF-7 cells were treated with vehicle (water) or abemaciclib (1μM for 8 days), replated and incubated with EdU (10 μM) for 20 hours, then stained with EdU/DAPI (scale bar, 100 μm) and quantified. n=3 independent experiments. **D** MCF-7 cells were treated with abemaciclib (1μM) for 30 days (refreshed every 48 hours), then the drug was washed out and cells were replated as single cells and cultured for an additional 30 days with drug-free media, white arrows indicate colonies formed by a single ‘escaped’ cell (scale bar, 1mm). **E** MCF-7 cells were treated with either vehicle (water) or abemaciclib (1μM for 8 days), then replated for treatment with common senolytic compounds. Cell viability was measured using MTS assay, n=3 independent experiments. For **A** and **B**, unpaired Student’s *t*-test (two-tail). For **E**, two-way ANOVA. Data are shown as mean ± SD. ***p<0.001.

### Abemaciclib induces lysosomal mass expansion in HR^+^ breast cancer

To uncover potential vulnerabilities induced by abemaciclib treatment, we analyzed the gene expression profile of treated and untreated MCF-7 cells using public RNA-seq data (Goel et al, 2017) in which a notable downregulation of cell cycle genes and upregulation of the senescence signature SenMayo were observed (Fig EV1B and C). Among the top differentially regulated genes, lysosomal genes and genes associated with lysosomal pathways and functions were significantly upregulated upon abemaciclib treatment at various concentrations and durations (Goel et al, 2017; Hafner et al, 2019; Watt et al, 2021) (Fig 2A and B and Fig EV2A-C). Elevated expression of lysosomal genes was validated by qPCR using an independent experimental sample set (Fig 2C). Transcription factor TFEB is a master regulator of lysosomal biogenesis. Interestingly, previous studies have demonstrated that CDK4/6 can phosphorylate TFEB and force its translocation from the nucleus to the cytoplasm, thereby inactivating its transcriptional function (Yin et al, 2020). In accordance, we observed that upon abemaciclib treatment TFEB was retained in the nucleus (Fig EV2D). We then evaluated how the altered expression of lysosomal genes impacted lysosomal content and structure. Acridine orange staining demonstrated an increase in the quantity of red fluorescent acidic vesicles, indicative of elevated lysosomal mass (Fig 2D). This observation was further supported by elevated CytoFix™ Red lysosomal staining, indicative of increased lysosomal mass (Fig 2E). Immunostaining revealed elevated expression of the lysosomal structural protein LAMP1, highlighting an increased number of lysosomes and changes in lysosomal size (Fig 2F and Fig EV2E). We further confirmed the increased fluorescence intensity of acridine orange, CytoFix™ Red, and LAMP1 in cells exposed to abemaciclib using flow cytometry (Fig 2G-I). Importantly, changes in lysosomal activity (SA-β-Gal), mass (acridine orange), and gene expression were observed in multiple HR^+^ breast cancer cell lines after abemaciclib treatment (Fig 2J-L and Fig EV2F). Finally, we examined whether lysosomal alterations could be detected in breast cancer patients treated with CDK4/6i. To accomplish this, we performed LAMP1 and CD63 immunostaining of tumor biopsies collected from the same patients before and after treatment. Strikingly, immunostaining for the lysosomal proteins LAMP1 and CD63 significantly increased after exposure to CDK4/6i in all patients (Fig 2M and N). Taken together, these data suggest that treatment with CDK4/6i causes consistent and reproducible lysosomal alterations in human cancer cells and biopsies.

**Figure 2.**
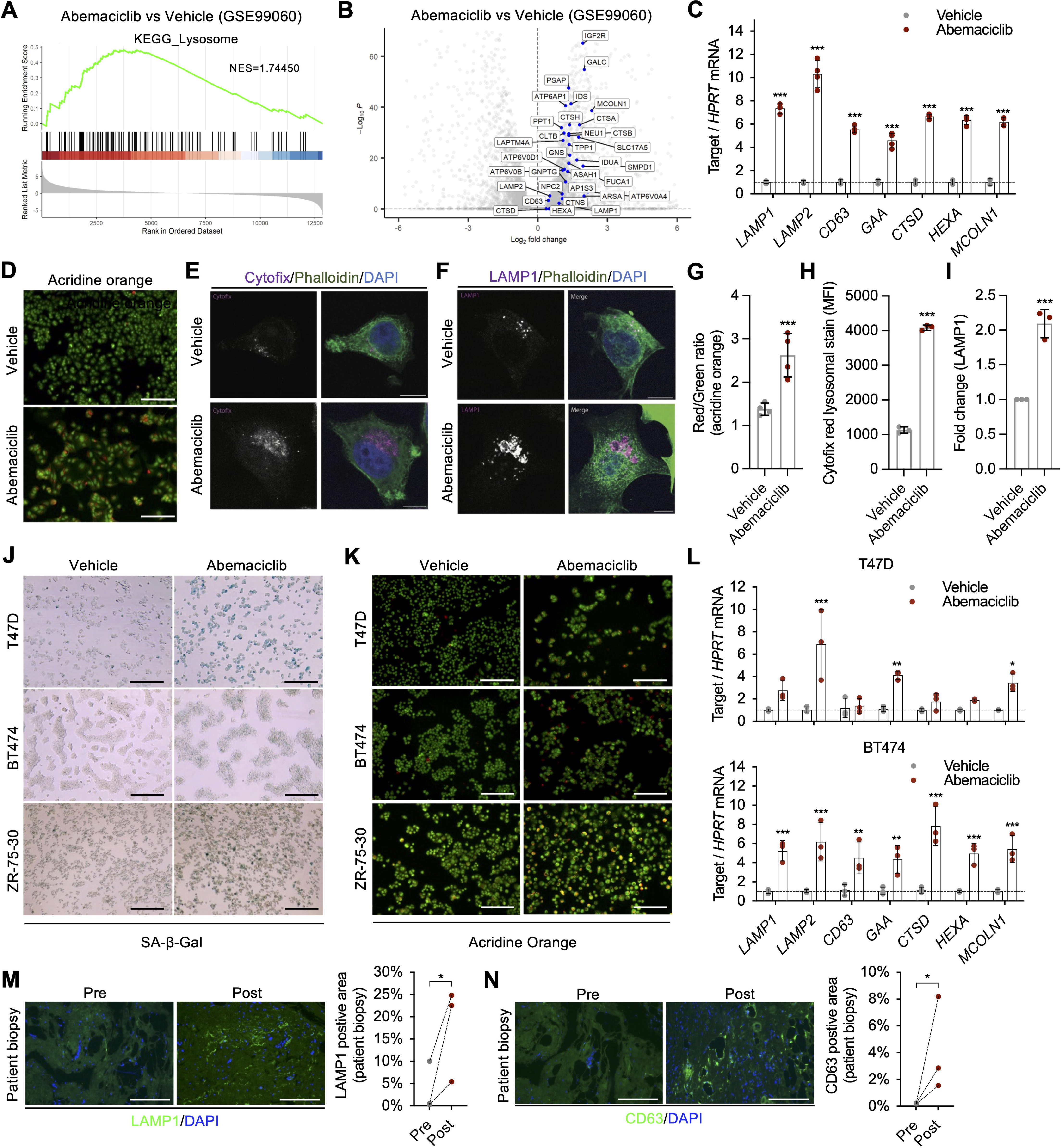
Abemaciclib increases lysosomal mass in HR+ breast cancer. **A,B** GSEA plot showing the enrichment for the KEGG_lysosome pathway geneset (**A**) and a volcano plot highlighting lysosomal genes that were upregulated (**B**) from abemaciclib treated MCF-7 cells compared to vehicle control. Expression data were obtained from GSE99060. **C-I** MCF-7 cells were treated with vehicle (water) or abemaciclib (1μM for 8 days), then cells were replated for qRT-PCR targeting lysosomal genes (**C**), acridine orange staining (scale bar, 1mm) (**D**), cytofix staining (scale bar, 10μm) (**E**), and LAMP1 staining (scale bar, 10μm) (**F**). The staining for acridine orange (**G**), cytofix (**H**) and LAMP1 (**I**) was quantified by flow cytometry analyses, n=3 or 4 independent experiments. **J-L** T47D, BT474 and ZR-75-30 cells were treated with vehicle (water) or abemaciclib (1μM for 8 days), then replated for SA-β-Gal staining (scale bar, 1mm) (**J**), acridine orange staining (scale bar, 1mm) (**K**), and qRT-PCR of lysosomal genes (**L**), n=3 independent experiments. **M-N** tumor biopsies from patients pre and post-treatment with CDK4/6 inhibitors were stained for LAMP1 (**M**) or CD63 (**N**), quantified, and compared (pre *vs* post), n=3 patients. For **C** and **L**, two-way ANOVA. For **G**, **H** and **I**, unpaired Student’s *t*-test (two-tail). For **M** and **N**, paired Student’s *t*-test (one-tail). *p<0.05, **p<0.01, ***p<0.001.

### Abemaciclib increases HR^+^ breast cancer cell sensitivity to lysosomotropic agents-induced cell death

An increase in lysosomal biogenesis along with changes in lysosomal size and structure could potentially render abemaciclib-treated cancer cells vulnerable to lysosomotropic agent-induced cell death. First, we used L-leucyl-L-leucine methyl ester (LLOMe), a lysosomotropic detergent assembled by lysosomal enzymes into a condensation product (Thiele & Lipsky, 1990). MCF-7 cells were exposed to abemaciclib (or vehicle) for 8 days, followed by treatment with LLOMe for 48 hours. The viability of cells exposed to sequential abemaclicb/LLOMe treatment was severely affected (Fig 3A). To monitor cell death in real time, we transduced MCF-7 cells with a mCherry-tagged histone 2b (H2b) reporter and incubated the cells with a green fluorescent indicator (Celltox^TM^ Green) of plasma membrane integrity loss. Interestingly, abemaciclib- treated cells exhibited high sensitivity to cell death upon exposure to LLOMe, as measured by the high number of double-positive cells relative to vehicle-treated cells (Fig 3B and C). Only a partial rescue of cell viability was obtained when we incubated Abemaciclib and LLOMe-treated cells with the pan-caspase inhibitor Q-VD-OPh (QVD), suggesting that caspase activation is not the sole inducer of cell death in this context (Fig EV2G). Loss-of-function mutations in oncosuppressor retinoblastoma protein 1 (RB1) are associated with increased resistance to CDK4/6 inhibitors (Antonarelli et al, 2023). Interestingly, analysis of an RNAseq dataset of MCF-7 cells carrying a short hairpin against RB1 and treated with abemaciclib revealed decreased expression of lysosomal genes (Fig EV2C). We observed a similar downregulation of lysosomal genes in abemaciclib-treated MCF-7 cells carrying siRNAs against RB1, which also correlated with a reduced sensitivity to LLOMe (Fig. EV2H and I). These data suggest that RB1 plays a role in modulating enhanced lysosomal biogenesis and sensitivity to lysosomotropic enzymes in cells treated with abemaciclib. Although LLOMe is a well-known and potent lysosomal detergent, its application in translational settings remains limited (Kavcic et al, 2020). Therefore, we tested salinomycin, a compound that has recently been shown to accumulate iron in lysosomes, consequently increasing ROS and leading to LMP (Mai et al, 2017), as well as bafilomycin A1, which inhibits vacuolar H^+^-ATPase (V-ATPase), causing lysosomal deacidification. Salinomycin effectively induced cell death in the abemaciclib-treated cells (Fig 3D and E). In contrast, no effect was observed with bafilomycin A1 (Fig 3F), suggesting that the abemaciclib-treated cells were insensitive to lysosomal deacidification. Based on these data, we selected salinomycin for further investigation. Similar to what was observed for MCF-7 cells, HR^+^ cancer cell lines BT474, T47D, and ZR-75-30 showed increased sensitivity to salinomycin- induced cell death following abemaciclib (Fig 3G-I). To determine whether sequential treatment with abemaciclib/salinomycin affects the viability of normal cells, we initially evaluated its effect in BJ and IMR90 fibroblasts. However, normal cells tolerated the treatment well, suggesting selective toxicity in cancer cells (Fig EV3A). To further validate this selectivity, we performed co-culture experiments with fluorophore-labelled MCF-7 cells and BJ fibroblasts. Sequential treatment with abemaciclib/salinomycin induced the death of MCF7 cells without affecting normal cells (Fig EV3B and C).

**Figure 3.**
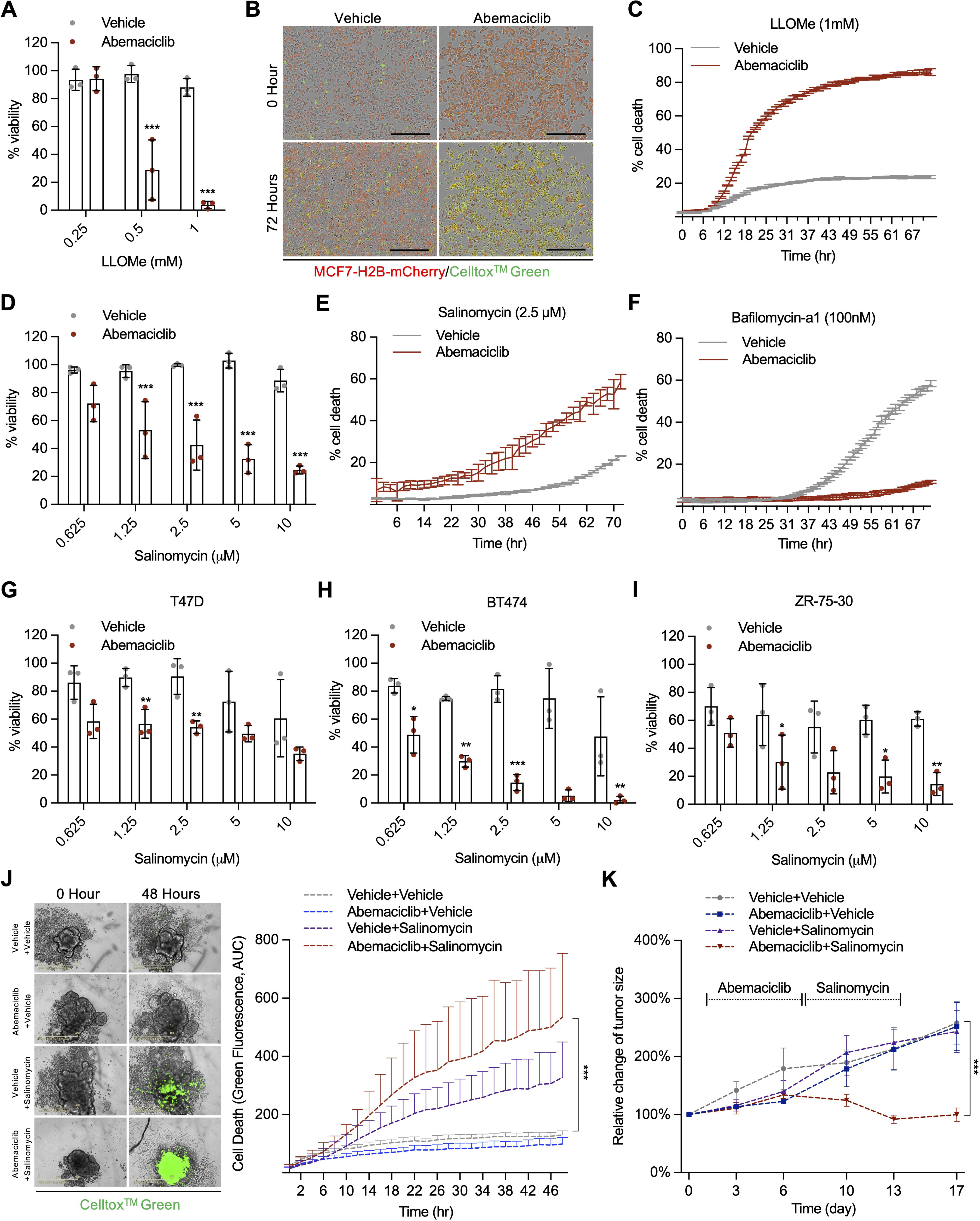
Abemaciclib sensitizes HR+ breast cancer cells to lysosomotropic agents-induced cell death. **A-F** MCF-7 cells (labeled with H2b-mCherry) were treated with vehicle (water) or abemaciclib (1μM for 8 days), then replated for subsequent treatment with LLOMe. Cell viability was measured by MTS assay (**A**) and cell death was assessed using IncuCyte live cell imaging with the death marker Celltox^TM^ Green, scale bar, 400μm (**B** for representative images and **C** for quantification). For subsequent treatments with salinomycin, cell viability was measured using MTS assay (**D**), while cell death was monitored using IncuCyte imaging (**E**). For treatment involving bafilomycin A1, cell death was measured through IncuCyte imaging (**F**). n=3 independent experiments. **G-I** Cells from T47D (**G**), BT474 (**H**), and ZR-75-30 (**I**) lines were treated with either vehicle (water) or abemaciclib (1μM for 8 days), then replated for further treatment with salinomyin. Cell viability was subsequently measured using MTS assay, n=3 independent experiments. **J** MCF-7 spheroids were formed, then treated with vehicle or abemaciclib (1μM for 6 days), followed by vehicle or salinomycin (5μM for 2 days) treatment. Cell death was measured using IncuCyte imaging with Celltox^TM^ Green and results were quantified and analyzed, n=4 spheroids/group. **K** Foxn1^Nu^ mice bearing MCF-7 tumors were treated with abemaciclib (40 mg/kg for 7 days) or salinomycin (5 mg/kg for 7 days) or both (abemaciclib/salinomycin sequential treatments). Tumor volume was measured over time and tumor growth curve was plotted, n=4 or 6 mice/group. For **A, C-I,** data are shown as mean ± SD, two-way ANOVA. For **J** and **K**, data are shown as mean ± SEM, two-way ANOVA. *p<0.05, **p<0.01, ***p<0.001.

Next, we evaluated whether sequential treatment could induce cancer cell toxicity in clinically relevant 3D structures. MCF-7 cells were grown in spheroids and exposed to abemaciclib and salinomycin, either alone or in combination. As observed in 2D cells, sequential treatment significantly increased the number of dead cells (Fig 3J). Finally, we tested the efficacy of the sequential treatment *in vivo*. MCF-7 cells were injected into the flanks of nude (*Foxn1*^Nu^) mice. The mice were then divided into different groups and treated with vehicle, abemaciclib, salinomycin, or a sequential combination of the two drugs. Compared with single treatment, sequential treatment effectively restrained tumor growth and decreased tumor size (Fig 3K). Altogether, these data suggest that sequential treatment with abemaciclib and lysosomotropic agents selectively achieves toxicity in HR^+^ breast cancer cells both *ex vivo* and *in vivo*.

### Abemaciclib induces a senescence-like phenotype in TNBC and enhances susceptibility to lysosomotropic agents

CDK4/6i, including abemaciclib, have received FDA approval for the treatment of advanced HR^+^, HER2^-^ metastatic breast cancer but are also under investigation in multiple pre-clinical and clinical studies for the treatment of other solid tumors, including triple-negative breast cancer (TNBC) (Hu et al, 2021). Most TNBC cells enter transient growth arrest during treatment with CDK4/6i in cell culture and quickly restore proliferation after drug withdrawal (Goel et al, 2017; Wang & Demaria, 2021). Accordingly, we observed that abemaciclib induced cell cycle arrest in the TNBC cell line MDA-MB-231 during treatment, with cells resuming proliferation upon drug withdrawal (Fig EV4A and B). Despite the lack of stable cell cycle arrest, abemaciclib-treated MDA-MB-231 cells showed alterations in lysosomal function, structure, and morphology, as exemplified by high SA-β-Gal activity, elevated expression of numerous lysosomal genes, and increased acridine orange and LAMP1 staining, reflecting altered lysosomal content and size (Fig 4A-F and Fig EV4C and D). In contrast, abemaciclib failed to promote lysosomal alterations in another TNBC cell line, BT549, which is resistant to CDK4/6i treatment (O’Brien et al, 2018). Interestingly, when we used a fluorescent probe to track CDK4/6 enzymatic activity, we observed high activity even in the presence of abemaciclib, suggesting alterations in the lysosomal biogenesis are consequence of the on-target effect of abemaciclib (Fig EV4E-I) Next, we evaluated whether abemaciclib rendered MDA-MB-231 cells more sensitive to the toxic effect of lysosomotropic agents. Similar to what was observed in HR^+^ cell lines, LLOMe and salinomycin, but not bafilomycin A1, showed significant toxicity in abemaciclib-treated MDA- MB-231 cells (Fig 4G and H and Fig EV4J), whereas no effect was observed in BT549 cells (Fig EV4K). Sequential treatment with abemaciclib and salinomycin effectively induced cell death in MDA-MB-231 3D spheroid cultures (Fig 4I). Finally, we tested sequential treatment *in vivo*. MDA-MB-231 cells were inoculated into the mammary fat pads of nude mice. Mice were then exposed to vehicle, abemaciclib, and salinomycin, either alone or in combination. Only for sequential treatment, a significantly reduced tumor size during the treatment phase and the absence of tumor growth after treatment withdrawal were observed (Fig 4J and K). These data suggest that sequential therapy can also be applied to TNBCs, provided that the activity of CDK4/6 is inhibited by abemaciclib.

**Figure 4.**
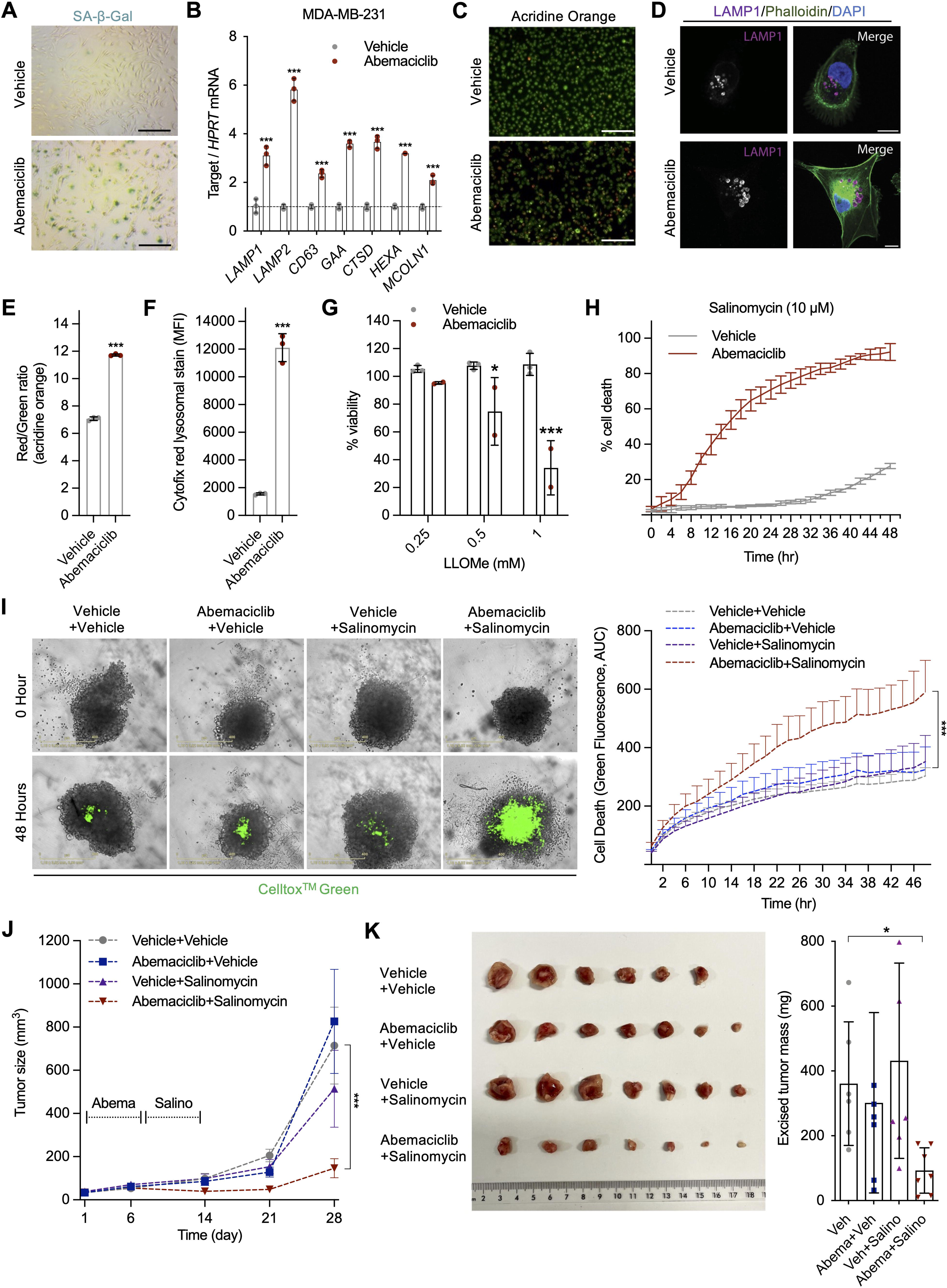
Abemaciclib sensitizes triple-negative breast cancer to lysosomotropic agents- induced cell death. **A-F** MDA-231 cells were treated with either vehicle (water) or abemaciclib (1μM for 8 days). After treatment, cells were replated, for SA-β-Gal staining (scale bar, 1mm) (**A**), qRT-PCR of lysosomal genes (**B**), acridine orange staining (scale bar, 1mm) (**C**), and LAMP1 staining (scale bar, 10μm) (**D**). The intensities of acridine orange (**E**) and cytofix (**F**) staining were quantitatively analyzed using flow cytometry. n=3 or 4 independent experiments. **G-H** Vehicle or abemaciclib (1μM for 8 days) treated MDA-231 cells were subsequently treated with LLOMe, and cell viability was quantified using MTS assay (**G**), or with salinomycin and their death was quantified using IncuCyte imaging with Celltox^TM^ Green (**H**), n=3 independent experiments. **I** MDA-MB-231 spheroids were treated with either vehicle or abemaciclib (1μM for 6 days) followed by vehicle or salinomycin (5μM for 2 days). Cell death was measured using IncuCyte imaging with Celltox^TM^ Green and quantification was plotted, n=4 spheroids/group. **J-K** Foxn1^Nu^ mice bearing MDA-231 tumors were treated with abemaciclib (40 mg/kg for 7 days) or salinomycin (5 mg/kg for 7 days) or both (abemaciclib/salinomycin sequential treatments), tumor volume was measured over time and data were used to plot tumor growth curve (**J**), excised tumors were weighed, and weights were plotted (**K**), n=6 or 7 mice/group. For **B, G** and **H,** data are shown as mean ± SD, two-way ANOVA. For **E** and **F**, data are shown as mean ± SD, unpaired Student’s *t*-test (two-tail). For **I** and **J**, data are shown as mean ± SEM, two-way ANOVA. *p<0.05, **p<0.01, ***p<0.001.

### Lysosomal enlargement enhances sensitivity to lysosomotropic agents-induced cell death

To assess the efficacy of other CDK4/6i in sensitizing breast cancer cells to lysosomotropic agents, MCF-7 cells were treated with palbociclib instead of abemaciclib. Interestingly, while palbociclib increased the sensitivity to both LLOMe and salinomycin, it did so to a lesser extent than abemaciclib (Fig 5A). Previous reports have demonstrated that abemaciclib, unlike other CDK4/6i inhibitors, induces intracellular vacuoles that originate from acidic vesicles, including late endosomes and lysosomes (Hino et al, 2020). We observed vacuoles in cells treated with abemaciclib, but not with palbociclib (Fig 5B). Consequently, abemaciclib-treated cells had larger lysosomes than palbociclib-treated cells, as revealed by LAMP1 staining (Fig 5C).

**Figure 5.**
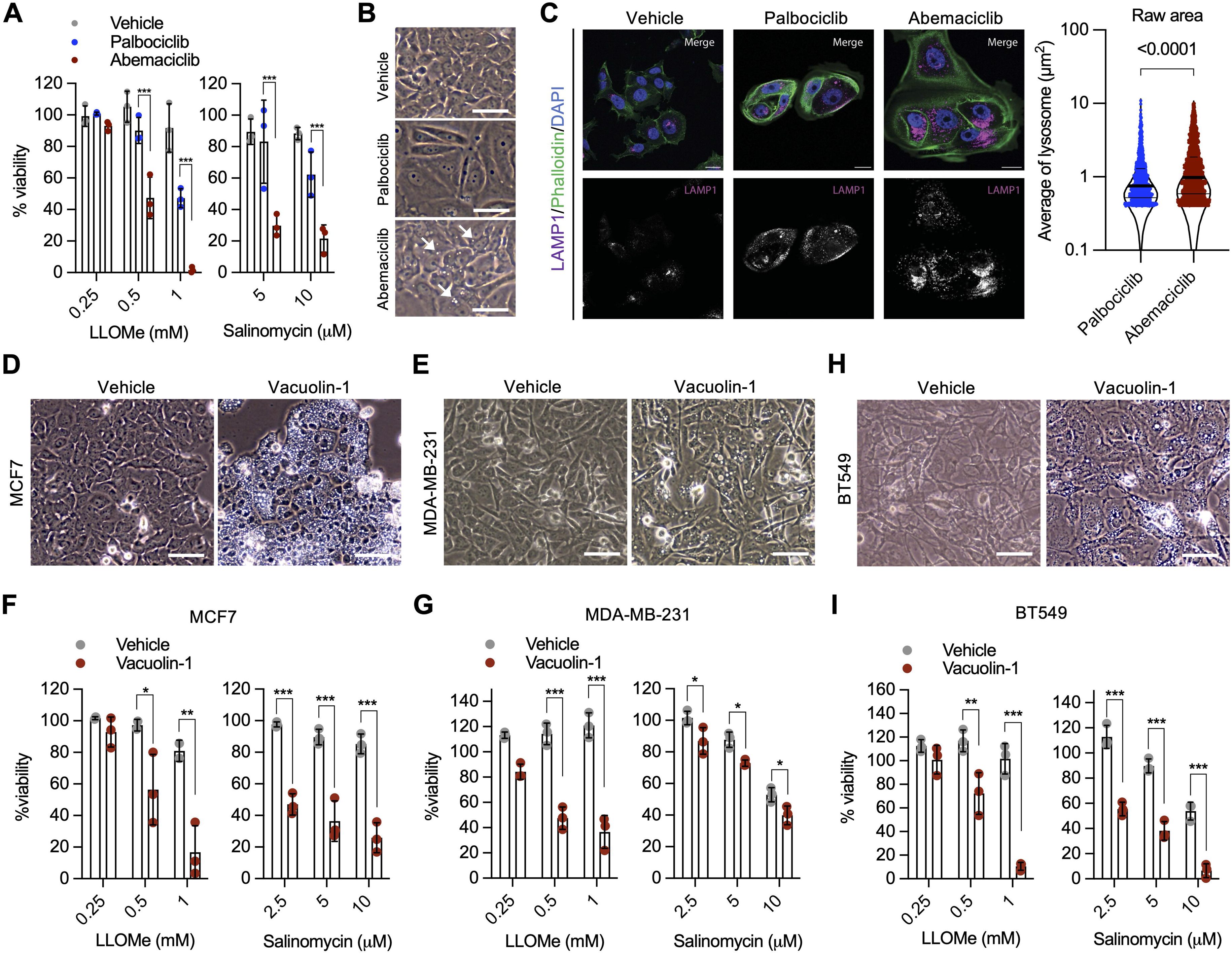
Lysosomal enlargement enhances sensitivity to lysosomotropic agents-induced cell death. **A-C** MCF-7 cells were treated with vehicle (water), palbociclib (1μM for 8 days), or abemaciclib (1μM for 8 days), then replated and subsequently treated with LLOMe or salinomycin. Cell viability was assessed using MTS assay (**A**). Cells were also imaged to identify vacuoles (white arrow; scale bar, 200μm) (**B**), and stained with LAMP1 (scale bar, 10μm), lysosomal size was quantified (**C**). n=3 independent experiments. **D-G** Cells were treated with vehicle or vacuolin-1, then imaged to visualize vacuoles formation (**D** for MCF-7 and **E** for MDA-MB-231; scale bar, 200μm), or subsequently treated with LLOMe or salinomycin and cell viability was quantified using MTS assay (**F** for MCF-7 and **G** for MDA-MB-231), n=3 independent experiments. **H-I** BT549 cells were treated with vehicle or vacuolin-1, then imaged for vacuoles visualization (**H**), or subsequently treated with LLOMe or salinomycin and cell viability was evaluated using MTS assay (**I**), n=3 independent experiments. Data are shown as mean ± SD, two-way ANOVA. *p<0.05, **p<0.01, ***p<0.001.

We then tested whether forcing the formation of vacuoles was sufficient to sensitize the cells to lysosomotropic agent-induced cell death. Vacuolin-1 is a cell-permeable compound that inhibits lysosomal fission and significantly increases the size of endo/lysosomal compartments, leading to vacuolization (Choy et al, 2018; Sano et al, 2016). Treatment of MCF-7 and MDA-MB-231 cells with vacuolin-1 resulted in a substantial increase in vacuole formation, but did not trigger a senescent phenotype (Fig 5D and E and Fig EV5A and B). Importantly, both MCF-7 and MDA- MB-231 cells exposed to vacuolin-1 were highly sensitive to LLOMe and salinomycin (Fig 5F and G). Finally, we evaluated whether vacuolin-1 could promote lysosomal alterations and vulnerability to lysosomotropic agents in cells insensitive to abemaciclib-induced lysosomal perturbations. Remarkably, the treatment of BT549 cells with vacuolin-1 resulted in pronounced intracellular vacuolization (Fig 5H) and enhanced sensitivity to lysosomotropic agent-induced cell death (Fig 5I). These findings indicate that the observed differences in sensitivity can be related, at least in part, to differences in lysosomal vacuolization, thereby extending the potential application of an alternative treatment approach to cancer cells that exhibit a limited response to CDK4/6i treatment.

## Discussion

CDK4/6 inhibitors, particularly abemaciclib, have emerged as promising agents for the treatment of HR^+^ and HER2^−^ breast cancer. However, several limitations of their efficacy remain, necessitating the development of improved therapeutic approaches. Prolonged inhibition of CDK4/6 activity in normal cells can promote various senescence-associated features, including durable and stable cell cycle arrest (Wang et al, 2022). Although cell cycle arrest is an intrinsic tumor suppressor mechanism, other senescence-associated phenotypic changes, including resistance to apoptosis and stemness, can contribute to breast cancer survival and dissemination (Evangelou et al, 2023). Thus, senescent-like cancer cells that can escape cell cycle arrest due to acquired genetic and epigenetic aberrations may have increased aggressiveness (McGrath et al, 2024). Therefore, sequential therapy based on a senescence-like phenotype inducer followed by a compound that takes advantage of the vulnerability developed by the initial treatment might be an efficient way to selectively eliminate tumor cells and reduce the risk of recurrence (Sieben et al, 2018). In this study, we showed that common senolytic compounds, most of which promote apoptosis, failed to eliminate breast cancer cells treated with abemaciclib. A particularly conserved phenotype across various models of senescence is the altered lysosomal activity and mass. Various studies have highlighted the significance of lysosomal biogenesis in the development of the senescence phenotype as well as its role in conferring resistance to different treatments (Curnock et al, 2023; Fassl et al, 2020; Li et al, 2023; Llanos et al, 2019; Martinez- Carreres et al, 2019; Rovira et al, 2022). Accordingly, we observed profound lysosomal changes in both HR^+^ and triple-negative breast cancer cells, which were independent of proliferative status. Importantly, analysis of breast cancer biopsies collected from human patients before and after treatment showed a CDK4/6i-induced increase in lysosomal mass. Additionally, we observed an enlargement in the lysosomal size. Lysosomal enlargement has been suggested to increase the sensitivity to lysosomotropic agent-induced cell death through lysosomal membrane permeabilization (Wang et al, 2018). We showed that a sequential therapeutic approach involving abemaciclib followed by lysosomotropic agents such as LLOMe and salinomycin was effective in inducing cell death in different HR^+^ breast cancer cells. Due to the fact that RB1 integrity is a common predictor of abemaciclib sensitivity, we investigated its role in lysosomal changes. We demonstrated that RB1 knockdown in abemaciclib-treated breast cancer cells resulted in reduced lysosomal biogenesis, accompanied by decreased sensitivity to lysosomotropic agent-induced cell death triggered by abemaciclib. These observed effects on lysosomal biogenesis may be directly or indirectly linked to RB activity. Suppression of RB can inhibit lysosomal biogenesis at the RNA level through other downstream factors. For instance, CDK1 suppression of TFEB activity suggests that active cell cycling can inhibit TFEB (Odle et al, 2020). However, other mechanisms that underlie this phenomenon cannot be excluded. In addition to HR^+^ breast cancer cells, abemaciclib treatment significantly increased lysosomal content and sensitivity to lysosomotropic agents in MDA-MB-231 cells, a TNBC cell line that is responsive to CDK4/6 inhibition. Conversely, BT549 cells demonstrated resistance to both abemaciclib and the subsequent lysosomotropic agent treatment. These findings highlight the heterogeneity within TNBC and underscore the need for precision medicine approaches that consider the specific cellular and molecular characteristics of different tumor types.

Previous studies have shown that in addition to general lysosomal enlargement, abemaciclib promotes aberrant lysosomal vacuolization (Hino et al, 2020). Interestingly, our data indicated that vacuolization was sufficient and necessary to sensitize breast cancer cells to the toxic effects of lysosomotropic agents, suggesting the potential for the development of more precise sequential therapeutic approaches that combine drugs promoting vacuolization with lysosomotropic agents. Therefore, identification of markers that can predict treatment efficacy is critical. It is logical to assume that an increase in lysosomal biogenesis and, potentially, lysosomal vacuolization may predict an increased sensitivity to cell death by compounds that can cause LMP. However, lysosomotropic agents induce LMP through different mechanisms of action, which rely on additional changes that follow the first treatment. Thus, further studies are needed to evaluate the involvement of different mechanisms that sensitize cancer cells to a specific type of treatment, allowing for a more refined prediction of treatment efficacy.

Overall, the sequential combinatorial strategy presented here supports the enormous potential of targeting aberrant lysosomal biology for the treatment of breast cancer. This strategy is distinct from traditional approaches targeting DNA or broad cellular pathways and offers an innovative solution to overcome the limitations of CDK4/6 inhibitor monotherapy and improve treatment outcomes.

## Materials and methods

### Cell culture and treatments

MCF-7, BT474, T47D, ZR-75-30, MDA-MB-231, BT549, and BJ cells were purchased from ATCC. All cells were cultured in DMEM-glutaMAX pyruvate medium (Thermo Fisher) supplemented with 10% fetal bovine serum (Thermo Fisher) and 1% penicillin-streptomycin (Lonza). Cells were kept in 5% O2, 5% CO2, 90% N2, and 37 °C incubators and were regularly tested for mycoplasma contamination.

For treatment, abemaciclib (MedCchemExpress, HY-16297) was dissolved in water and diluted in DMEM media to a final concentration of 1uM. The lysosomotropic agents L-Leucyl-L- Leucine methyl ester (Cayman, 16008) and salinomycin (MedChemExpress, HY-15597) were dissolved in DMSO. Bafilomycin-A1 (MedChemExpress, HY-100558) and Q-VD-OPH (MedChemExpress, HY-12305) were also dissolved in DMSO. All drugs were further diluted in DMEM media to treat cells at different concentrations, as indicated in each figure. Senolytic drugs were dissolved in DMSO and further diluted in DMEM media to treat cells at different concentrations, as indicated in Figure 1: Navitoclax (ABT-263) (MedChemExpress, HY-10087), Dasatinib hydrochloride (MedChemExpress, HY-10181A), Quercetin hydrate (Sigma-Aldrich, 337951), Fisetin (MedChemExpress, HY-N0182), Nutlin-3a (MedChemExpress, HY-10029), Piperlongumine (MedChemExpress, HY-N2329), Geldanamycin (HSP90 inhibitor) (Cayman Chemical, 13355).

### Colony formation assay

Drug-treated breast cancer cells were replated in a 6-well plate (5 × 10^3^ cells/well) at the end of treatment and allowed to grow in drug-free normal medium for 8 days. Afterward, cells were fixed in 4% PFA for 30 min and then stained with 0.2% crystal violet in 37% methanol for 1 hour. Pictures of all the plates were taken with a scanner (Epson). The images were cropped and processed in Microsoft PowerPoint using the same settings.

### EdU staining

Control and treated cells were replated on coverslip in a 24-well plate (3 × 104 cells/well) and cultured for 20 hours in the presence of EdU (10 μM), then fixed and stained as previously described (Kohli et al, 2021). Images were acquired at 100 times magnification (Leica), and the number of cells was counted with ImageJ software.

### Senescence-Associated β-Galactosidase staining

Control and treated cells were re-plated in 24 well plate. 24 hours later, the cells were fixed and incubated with Sa-β-galactosidase staining, as previously described (Kohli et al, 2021). Images were acquired using a Leica microscope and processed using the ImageJ software.

### MTS viability assay

The cells were seeded in 96-well plates. Drug treatments were performed directly on the plate and three technical replicates were used for each experimental condition. After 24 or 48 h of treatment, cell viability was measured using MTS solution (CellTiter 96® Aqueous Non- Radiative Cell Proliferation Assay, Promega) mixed with the culture medium following the manufacturer’s instructions. Absorbance was recorded at 450 nm and normalized to the reference absorbance measured in the acellular wells.

### Lentivirus production and transduction

For the production of lentiviral particles, 293FT cells were plated in a 10 cm Petri dish. 24 hours later, cells were transfected with a mix of plasmids encoding viral components (ViraPOwer plasmid mix, K497-00, Invitrogen) in addition to the lentiviral plasmid of interest using TurboFect™ Transfection Reagent (R0532, Thermo Fisher) overnight. The next day, the transfection mix was replaced with normal growth media, collected 24 or 48 h later, and used immediately for transducing cells or frozen at -80 °C. The pLenti6-H2b-mCherry plasmid was a gift from Torsten Wittmann (Addgene, plasmid # 89766), pLenti-mCherry-CDK4KTR plasmid was a gift from Hee Won Yang (Addgene, plasmid #126680). Lentiviral particles harbor the H2b-mcherry construct were used to generate cells with stable nuclear mCherry. Lentiviral particles harbor the mCherry-CDK4KTR construct were used to generate cells with stable CDK4/6 kinase activity reporter. For transduction, cells were plated in 6 well plate and once attached, viral particles were added in addition to normal growth media, and then the plate was centrifuged at 4000xg for 40 min. The next day, the cells were washed and refreshed with complete medium, and two days later, the cells were selected with blasticidin (2.5 uM).

### Real-time imaging of cell proliferation and death

Time-lapse imaging was performed using the IncuCyte® ZOOM system to examine cell death and proliferation in real-time. Cells stably expressing a nuclear-localized red fluorescent (H2b- mcherry-labelled), were cultured in the presence of Celltox green, a non-toxic cell-impermeable dye that emits green fluorescence once it binds to nucleic acids. Therefore, once the plasma membrane is permeabilized, the dye can enter the cell and bind to nuclear DNA. Double-positive cells (green and red) were considered dead. The number of red and double-positive cells was automatically quantified using IncuCyte ZOOM software.

### Acridine orange staining

For Acridine Orange staining, the cells were first washed and then incubated with 1 μM acridine orange (Sigma-Aldrich) in PBS for approximately 20 min at 37°C. The cells were then rinsed and stored in complete medium or PBS during image acquisition. Images were acquired in red and green channels using a Leica microscope. Images were processed using the ImageJ software.

### LAMP1 cellular staining

Immunofluorescence for LAMP1 was conducted by seeding cells on a glass cover slip (Aurion 72231-01) at 60% cell density after senescence induction. Cells were fixed with 4% PFA and washed three times with PBS followed by a blocking step with 0.2-micron filtered 3% BSA and 0.15% glycine in PBS which was supplemented with 0.1% saponin for 30 min at room temperature. Next, samples were stained with LAMP1 Ab (Bioke, 9091S) overnight at 4 degrees, followed by three washes with PBS and consecutive secondary staining with anti-rabbit- AlexaFluor633 (Thermo Fisher Scientific, A-21072), phalloidin-AlexaFluor488 (AAT Bioquest, 23115) and DAPI (Sigma-Aldrich, D5942-5MG). After the final three PBS washes to get rid of unbound reagents, samples were mounted on glass microscopy slides and imaged using SP8(.X) with a 63X magnification with a N/A of 1.4. image analysis was conducted with an imageJ script (see supplementary information) with further processing of the data using pandas in Pyhthon.

### Flow cytometry

Cells were incubated with 1µg/ml Acridine Orange (Sigma-Aldrich, 235474-5G) or a 1:1000 dilution of pre-dissolved CytoFix™ Red Lysosomal Stain (AAT Bioquest, 23210) in DMEM with 10% FBS for 30 min at 37C and 5% CO2. Next cells were trypsinized from the plate and washed twice in cold PBS with 1% BSA before measuring the samples on the Canto II. Flow cytometry analysis was conducted with FlowJo V10. For the LAMP1 staining, cells were fixed after being trypsinized from the plate and fixated with 4% PFA followed by a protocol that was described in the section ’Lysosomal analysis by microscopy’, but only using the anti-rabbit- AlexaFluor633 as secondary.

### Bioinformatics

We performed the gene expression analysis using the ’DESeq2’ package (Love et al, 2014), applying an alpha cutoff of 0.05 and using the Bonferroni p-value adjustment method. We performed Gene Set Enrichment Analysis (GSEA) based on the fold-change values obtained from DESeq2 and visualized the results using the ’gseGO’ function from the ’clusterProfiler’ package (Wu et al, 2021). We generated the volcano plot using the ’EnhancedVolcano’ package. All RNA-seq data were analyzed using R version 4.2.3.

### Real-time PCR

An Isolate II RNA Mini Kit (Bioline, Cat# BIO-52073) was used for total RNA isolation. For reverse transcription, 500ng of RNA was transcribed into cDNA using a kit (Applied Biosystems, Cat# 4368813). qRT-PCR reactions were performed using GoTaq® qPCR Master Mix (A6002, Promega). Tubulin was used as a housekeeping gene to normalize the expression of target genes. List of primers:

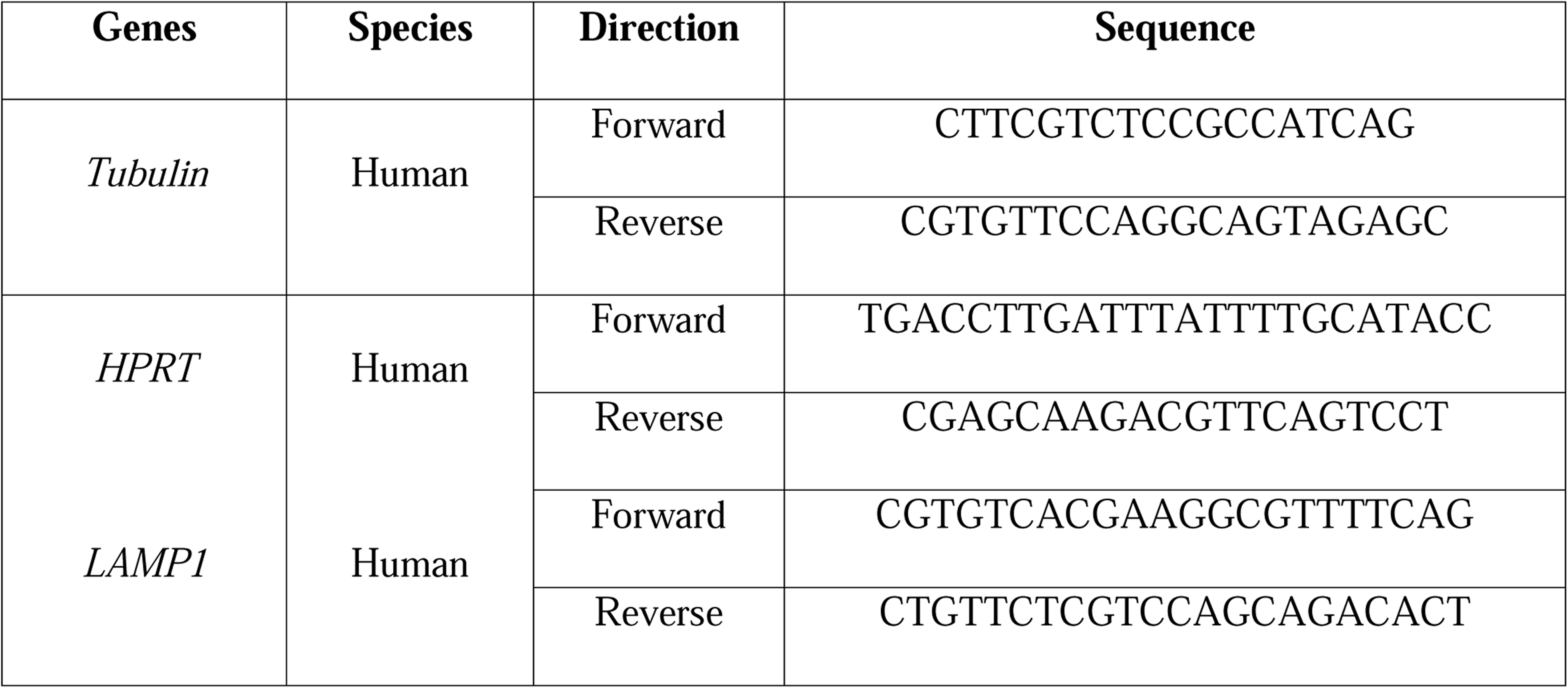
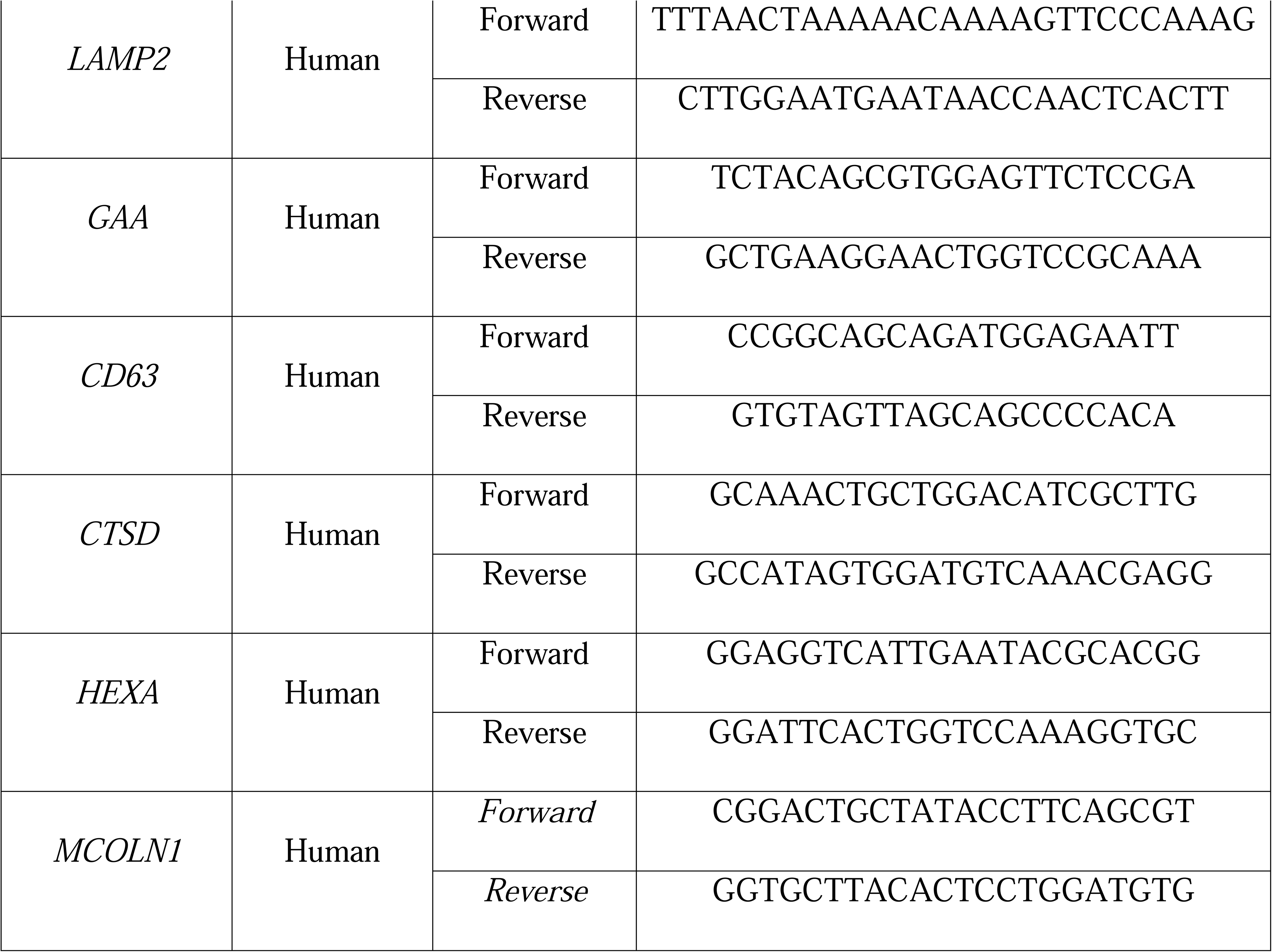

### Spheroids

Breast cancer spheroids were constructed as described before(Soto-Gamez et al, 2022). Briefly, cells were seeded at a density of 1500 cells per well in ultra-low attachment 96-well plates (Corning Incorporated, Kennebunk, ME, USA). Plates were centrifuged at 1000 rpm for 5 min to promote cell aggregation and spheroid formation. After 6 days of incubation, spheroids were suitable for performing experiments and cell senescence was induced by treatment with abemaciclib (1 µM) for 6 additional days. Spheroids were next treated with salinomycin (5 µM) or vehicle (DMSO) for 48 hours. Cell death was monitored in a time-resolved assay, using CellTox™ Green Cytotoxicity Assay (Promega) to measure fluorescent increases every 2 hours with an IncuCyte S3 (Essen BioScience). The mean green fluorescence intensity relative to the initial time point (0h00m), and area under the curve (AUC) for each of the conditions tested was calculated.

### Mouse xenografts

All mice were maintained in individually ventilated cages (IVC) at the Central Animal Facility (CDP) of the University Medical Center Groningen (UMCG) under standard conditions. The experiments were approved by the Central Authority for Scientific Procedures on Animals (CCD) of the Netherlands. 12-week-old female *Foxn1*^Nu^ (Jackson Laboratory) mice were used to generate the MCF-7 and MDA-MB-231 xenograft cancer models.

17β-estradiol pellets (0.18 mg for 90-day release, catalog number E-121, Innovative Research of America) were transplanted to the left flanks of 12-week-old female *Foxn1*^Nu^ mice, one week later, MCF-7 cells (2 × 10^6^) were injected into the right flanks. Approximately 3 weeks after injection, mice were randomly divided into four groups and injected with PBS or abemaciclib (40 mg/kg) for 7 consecutive days, followed by vehicle or salinomycin (5 mg/kg) for an additional 7 consecutive days. Tumor growth was calculated by measuring the tumor length and width using a calliper, and the investigator was blinded. Mice exhibiting skin side effects due to estrogen pellets were excluded from the experiment before the start of treatment injections.

MDA-231 cells (2 × 10^6^) were injected into the right mammary fat pad of 12-week-old female *Foxn1*^Nu^ mice. Approximately 3 weeks after injection, mice were divided into four groups and injected with PBS or abemaciclib (40 mg/kg) for 7 consecutive days, followed by vehicle or salinomycin (5 mg/kg) for an additional 7 consecutive days. Tumor growth was calculated by measuring the tumor length and width using a calliper, and the investigator was blinded. After excision, the weight of each tumor was measured. The weight of each mouse was measured once a week.

### Patients

Formalin fixed paraffin embedded tumor samples from three hormone receptor positive, HER2 negative metastatic breast cancer patients were collected at IRCCS Istituto Romagnolo per lo Studio dei Tumori (IRST) “Dino Amadori”, Meldola, Italy before and after exposure to treatment with CDK4/6 inhibitors (one treated with abemaciclib, one with ribociclib and one with palbociclib) and hormone therapy with aromatase inhibitors. All samples were collected according to a protocol approved by the Italian Local Ethics Committee (CEROM IRST IRCCS- AVR, protocol code: IRST B114) and all the patients have signed informed consent. The procedure was performed before starting the treatment and at the time of the best clinical response, respectively 6.9, 11.8 and 7.6 months after starting treatment. At the last follow-up, the three patients were disease-free. Samples were stained with LAMP1 primary antibody (9091S, Bioke) and CD63 primary antibody (NBP2-42225, Novus biologicals) overnight at 4 degrees, followed by three washes with PBS and a consecutive staining with secondary antibodies respectively and DAPI (Sigma-Aldrich, D5942-5MG). After the final three PBS washes to get rid of unbound reagents, samples were mounted and imaged at 100 times magnification (Leica).

## Declarations

### Ethical approval

The mouse experiments were approved by the Central Authority for Scientific Procedures on Animals (CCD) of the Netherlands. All human samples were collected according to a protocol approved by the Italian Local Ethics Committee (CEROM IRST IRCCS-AVR, protocol code: IRST B114) and all the patients have signed informed consent.

### Authors’ contributions

J.N., B.W. and M.D. conceptualized the study. J.N., S.M., A.S.-G., B.W., and M.D. devised the methodology. J.N., S.M., S.B., M.Z., C.G., U.D.G., A.S.-G., A.A., T.G., and B.W. carried out the investigation. J.N., S.M., A.A. and B.W. visualized the data. B.W. and M.D. supervised the project. J.N. and M.D. wrote the original paper draft. M.D. acquired the funding and managed the project. J.N., S.B., B.W. and M.D. reviewed and edited the paper.

### Availability of data and materials

Data is provided within the manuscript or supplementary information files. All the datasets and access numbers used for the study are cited in their relative descriptions. Additional reasonable requests can be made directly to the corresponding authors.

