## Supplementary Figures for "Pharmacological CDK4/6 inhibition promotes vulnerability to lysosomotropic agents in breast cancer"

### Expanded View Figures

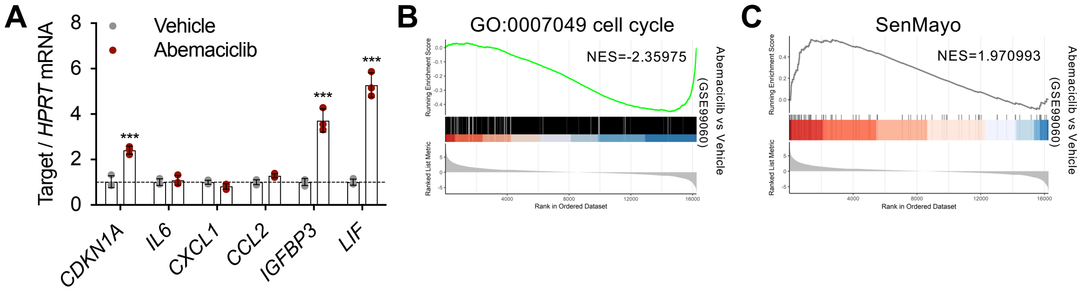


**Figure EV1 Abemaciclib treatment induces senescence-like phenotypes in MCF-7 breast cancer.**

**A** RNA was extracted from MCF-7 cells treated with either the vehicle (water) or abemaciclib (1 µM for 8 days), followed by qPCR analysis targeting the specified genes. n=3 independent experiments.

**B, C** Gene Set Enrichment Analysis (GSEA) plot showing the enrichment for the GO term “cell cycle” (**B**) and the SenMayo geneset (**C**) in MCF-7 cells treated with abemaciclib compared to vehicle-treated cells. Expression data were obtained from GSE99060.

Data are mean ± SD. For **A**, two-way ANOVA, ***p<0.001.

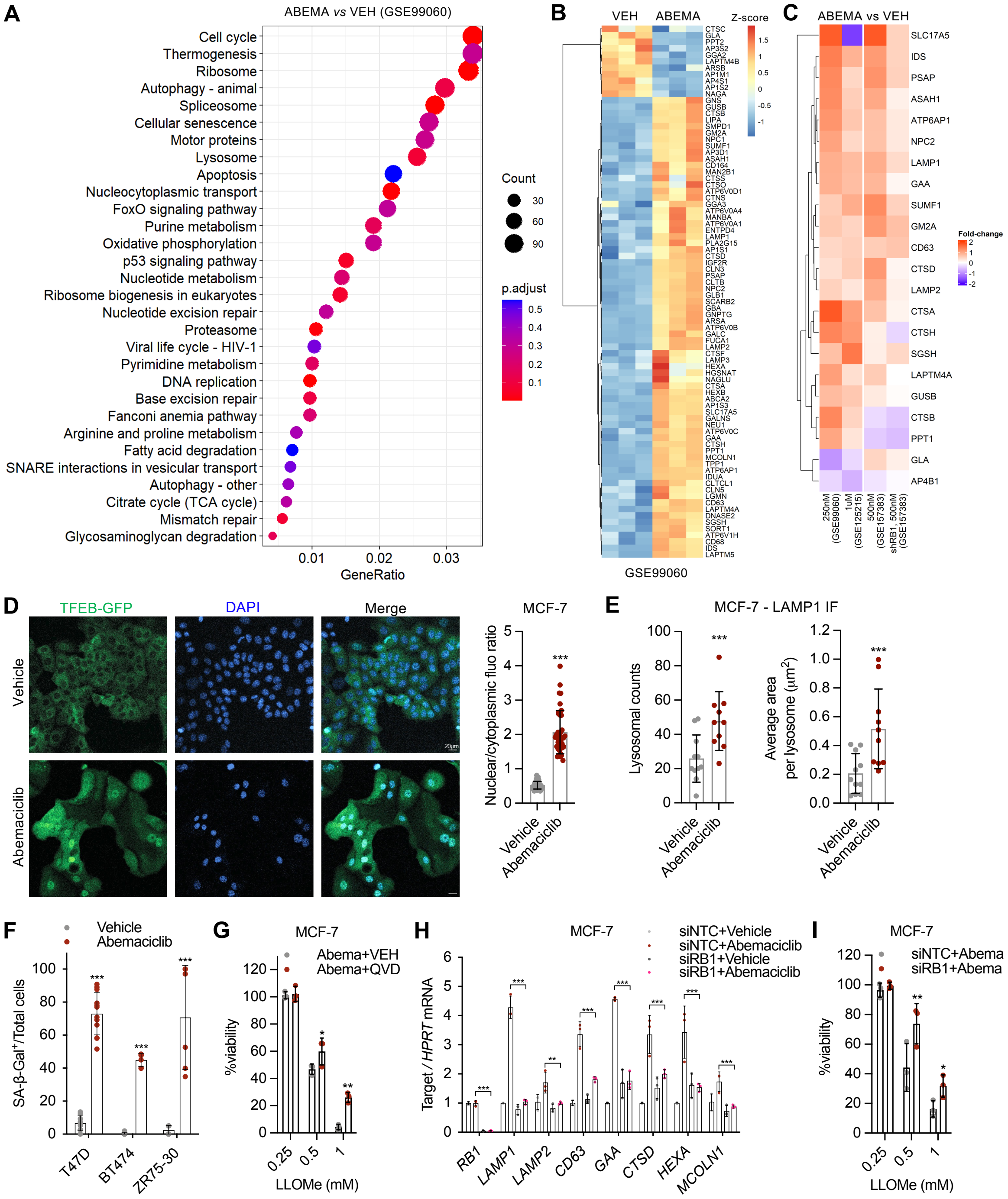


**Figure EV2 Abemaciclib treatment increases lysosomal mass in HR$^+$ breast cancer.**
**A** Dot plot shows the significantly enriched GO terms when comparing the gene expression profile of MCF-7 cells treated with abemaciclib to vehicle-treated cells.
**B** Heatmap of differentially expressed lysosomal genes following abemaciclib treatment in MCF-7 cells. Expression data were obtained from **GSE99060**.
**C** Heatmap of differentially expressed lysosomal genes following abemaciclib treatment (fold change *vs* vehicle) in MCF-7 cells with indicated doses and genetic backgrounds. Expression data were obtained from **GSE99060**, **GSE125215**, **GSE157383**.
**D** MCF-7 cells labeled with TFEB-GFP via lentiviral particles were treated with either the vehicle (water) or abemaciclib (1 µM for 48 hours), followed by imaging to assess the

subcellular localization of TFEB proteins (scale bar: 20 µm). The ratio of nuclear to cytoplasmic fluorescence mean signal was quantified and plotted. n=3 independent experiments.

**E** MCF-7 cells were treated with either the vehicle (water) or abemaciclib (1 µM for 8 days), replated, and stained for LAMP1. Cells were then analyzed to quantify lysosomal counts and the average lysosomal area. Cells are from 3 independent experiments.

**F** Quantification of SA-β-Gal positive cells from **Figure 2J**.

**G** MCF-7 cells were treated abemaciclib with or without QVD (5µM), followed by treatments with LLOMe at the indicated concentrations. Cell viability was measured using MTS assay. n=3 independent experiments.

**H** MCF-7 cells were transfected with siRNA targeting *RB1* and then treated with vehicle or abemaciclib (1µM for 5 days) starting 1 day after siRNA transfection. At the end of the treatment, RNA was extracted from both siNTC and siRB1 groups, and qPCR was performed to assess the expression of *RB1* and lysosomal genes. n=3 independent experiments.

**I** MCF-7 cells were transfected with siRNA targeting RB1 and treated with either the vehicle or abemaciclib (1 µM for 5 days), starting 1 day after siRNA transfection. Following this, the cells were replated and subjected to subsequent treatments with LLOMe for 48 hours at the indicated concentrations. n=3 independent experiments.

Data are mean ± SD. For **D** and **E**, unpaired Student's *t*-test (two-tail). For **F-I**, two-way ANOVA. *p<0.05, **p<0.01, ***p<0.001.

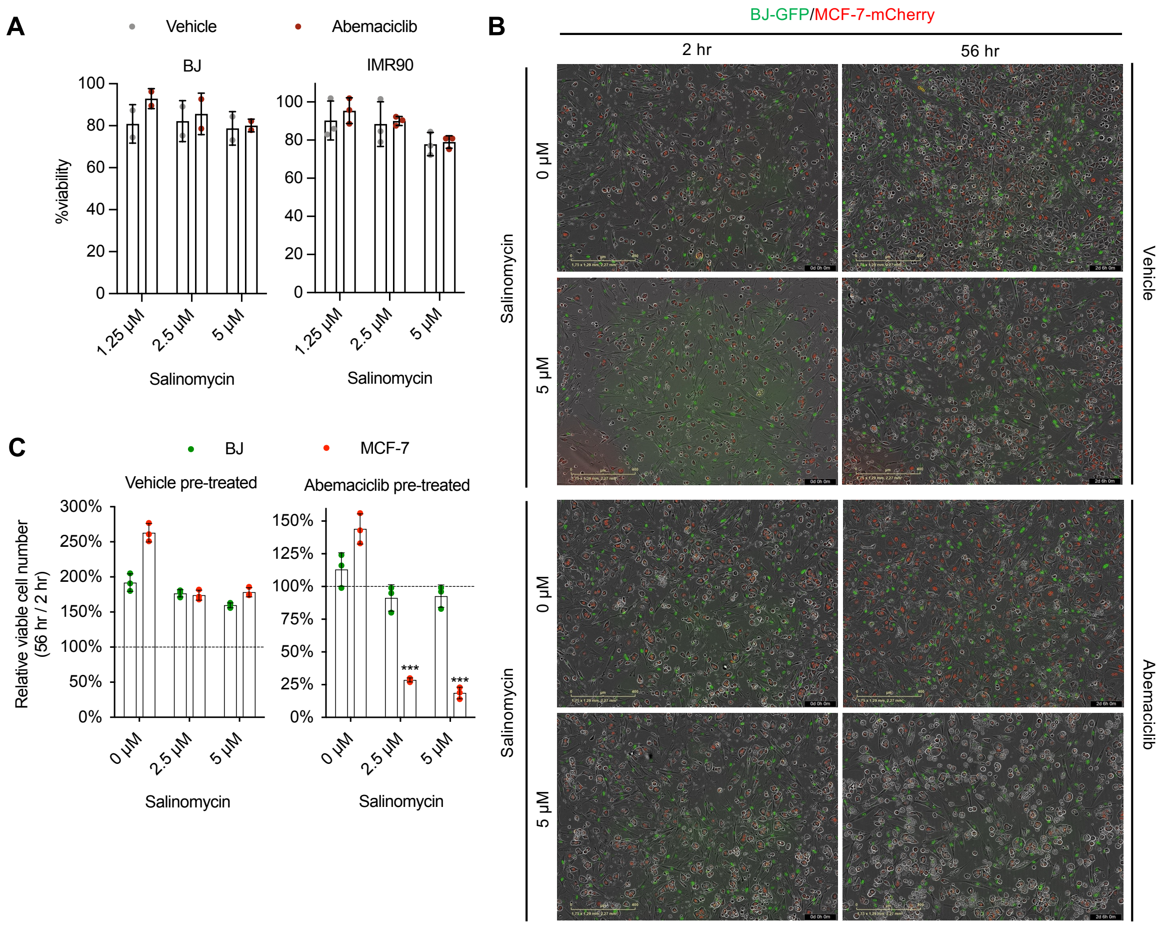


**Figure EV3 Abemaciclib-treated normal fibroblasts are not sensitive to lysosomotropic agents.**

**A** BJ or IMR90 cells were pretreated with either vehicle (water) or abemaciclib (1µM for 7 days), followed by incubation with salinomycin at the indicated concentrations. Cell viability was then assessed using an MTS assay. n=3 independent experiments.

**B** MCF-7 cells (mCherry) and BJ cells (GFP) were pretreated with either vehicle (water) or abemaciclib (1µM for 6 days) and then co-cultured. These cells were subsequently incubated with salinomycin (0 or 5µM for 56 hours), with representative images captured using IncuCyte at 2 and 56 hours.

**C** Ratio of viable MCF-7 and BJ cells (56-hour vs 2-hour) treated with different concentrations of salinomycin, for both vehicle and abemaciclib pretreated groups. n=3 independent experiments.

Data are mean ± SD. Two-way ANOVA. ***p<0.001.

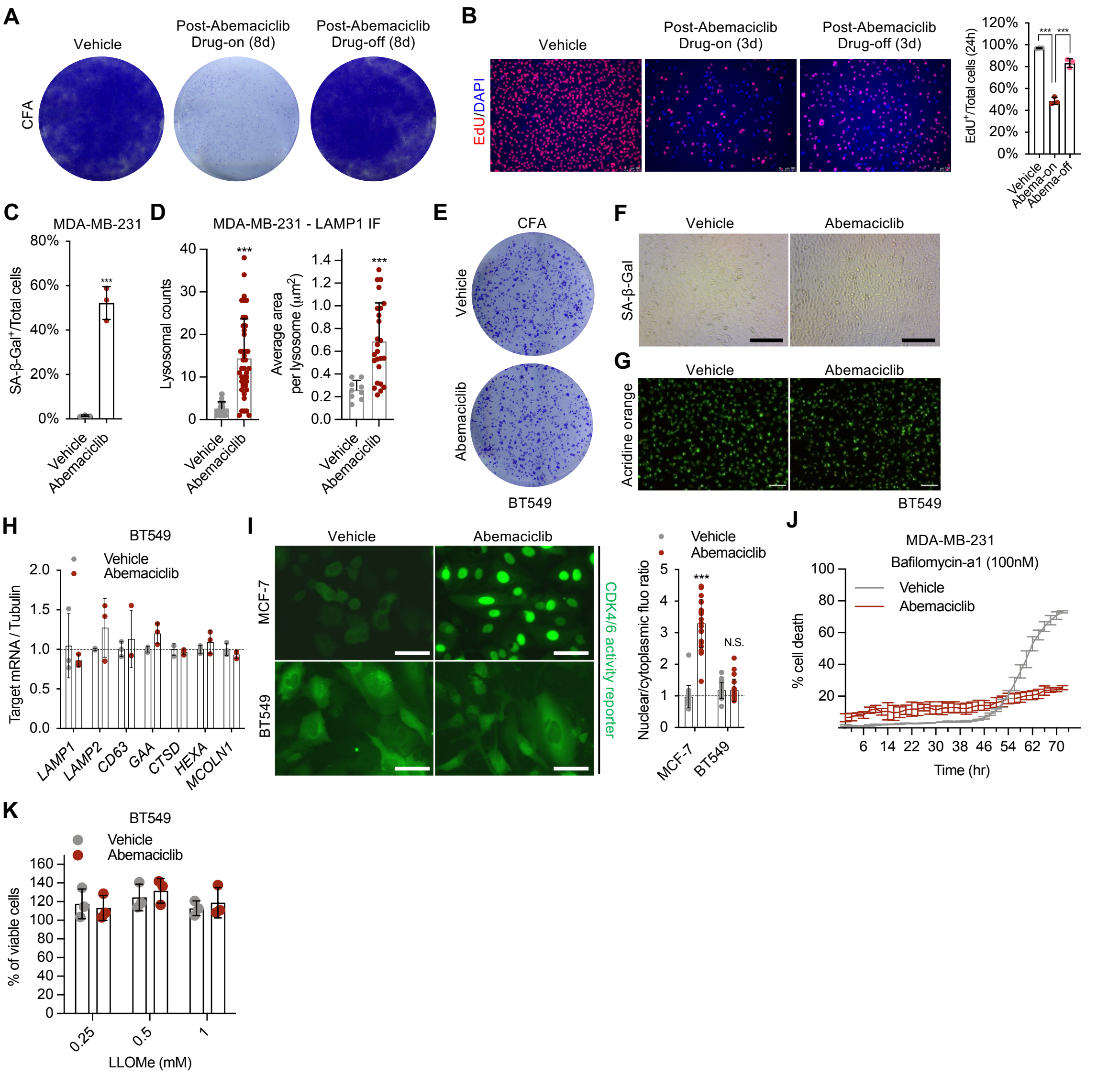


**Figure EV4 Abemaciclib selectively sensitizes triple-negative breast cancer cells to lysosomotropic agents-induced cell death**

**A-B** MDA-MB-231 cells were treated with either the vehicle (water) or abemaciclib (1 μM for 8 days) and then replated for subsequent assays. For the colony formation assay (**A**), cells were cultured with or without abemaciclib for an additional 8 days, followed by staining. For EdU staining (**B**), cells were replated and treated with or without abemaciclib for 3 days. EdU (10 μM) was added on day 4 for 20 hours of incubation, followed by staining (scale bar: 100 μm).

**C** Vehicle or abemaciclib (1μM for 8 days) treated MDA-MB-231 cells were stained with LAMP1 and analyzed for lysosomal counts and average area.

**D** MDA-MB-231 cells were treated with vehicle (water) or abemaciclib (1μM for 8 days), then replated and stained with LAMP1, and cells were quantified for lysosomal counts or average area of lysosomes. Cells are from 3 independent experiments.

**E-H** BT549 cells were treated with vehicle (water) or abemaciclib (1μM for 8 days), then cells were replated for colony formation assay (8 days culture) (**E**), SA-β-Gal staining (scale bar, 1mm) (**F**), acridine orange staining (scale bar, 1mm) (**G**), or qRT-PCR for lysosomal genes (**H**), n=3 independent experiments.

**I** MCF-7 or BT549 cells were transfected with lentiviral particles encoding a CDK4/6 kinase activity reporter (CDK4KTR), followed by treatment with either the vehicle or abemaciclib (1 μM for 48 hours). Cells were imaged to analyze the localization of mCherry protein (scale bar: 60 μm). Cytoplasmic localization of mCherry indicates active CDK4/6 kinases, while nuclear localization indicates suppressed CDK4/6 kinase activity. The nuclear-to-cytoplasmic mCherry fluorescence mean intensity ratio was calculated for each cell and plotted. Data represents cells from 3 independent experiments.
**J** Vehicle or abemaciclib (1μM for 8 days) pretreated MDA-MB-231 cells were subsequently treated with bafilomycin A1 and cell death was measured using IncuCyte live cell imaging with Celltox™ Green. n=3 independent experiments.
**K** Vehicle or abemaciclib pretreated BT549 cells were subsequently treated with LLOMe and viability was measured using MTS assay. n=3 independent experiments.
Data are mean ± SD. For **B**, one-way ANOVA. For **C** and **D**, unpaired Student's *t*-test (two-tail). For **H-K**, two-way ANOVA. ***p<0.001.

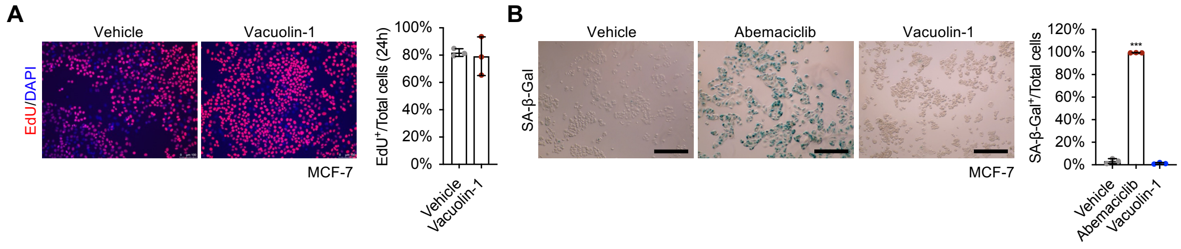


**Figure EV5 Vacuolin-1 is unable to induce senescence-like phenotype**

**A-B** MCF-7 cells were treated with vehicle or vacuolin-1 (1μM for 8 days) and replated for EdU staining (scale bar, 100μm) (**A**) or SA-β-Gal staining (scale bar, 1mm) (**B**). n=3 independent experiments.

Data are mean ± SD, one-way ANOVA. ***p<0.001.
